# Biological age is a universal marker of aging, stress, and frailty

**DOI:** 10.1101/578245

**Authors:** Timothy V. Pyrkov, Peter O. Fedichev

**Affiliations:** Gero LLC, 105064, Nizhny Susalny per. 5/4, Moscow, Russia; Moscow Institute of Physics and Technology, 141700, Institutskii per. 9, Dolgoprudny, Moscow Region, Russia

## Abstract

We carried out a systematic investigation of supervised learning techniques for biological age modeling. The biological aging acceleration is associated with the remaining health- and life-span. Artificial Deep Neural Networks (DNN) could be used to reduce the error of chronological age predictors, though often at the expense of the ability to distinguish health conditions. Mortality and morbidity hazards models based on survival follow-up data showed the best performance. Alternatively, logistic regression trained to identify chronic diseases was shown to be a good approximation of hazards models when data on survival follow-up times were unavailable. In all models, the biological aging acceleration was associated with disease burden in persons with diagnosed chronic age-related conditions. For healthy individuals, the same quantity was associated with molecular markers of inflammation (such as C-reactive protein), smoking, current physical, and mental health (including sleeping troubles, feeling tired or little interest in doing things). The biological age thus emerged as a universal biomarker of age, frailty and stress for applications involving large scale studies of the effects of longevity drugs on risks of diseases and quality of life.

To be published as Chapter 2 in “Biomarkers of aging”, ed. A. Moskalev, Springer, 2019.

## Background

Aging in most species, including humans, manifests itself as a progressive functional decline leading to increasing prevalence (Mitnitski and Rockwood 2016, Yu et al. 2017) and incidence of the chronic age-related diseases, such as cancers, diabetes, cardiovascular diseases (Niccoli and Partridge 2012, Podolskiy et al. 2016, Zenin et al. 2019) or disease-specific mortality (Barzilai and Rennert 2012), all accelerating exponentially at a rate compatible with that of the Gompertz mortality law (Gompertz 1820, Makeham 1860). The physiological indices or physiological state variables, such as blood biochemistry or cell count markers change with age in a highly coordinated manner. This can be best seen from a Principal Component Analysis (PCA), which is a mathematical procedure aimed at the identification of the most correlated or collective features in multidimensional data (Ringnér 2008). The method is an almost always the first choice in biological data analysis and searches for aging signatures in a multitude of biological signals ranging from blood samples to locomotor activity patterns (Pyrkov et al. 2018b), anthropological and functional measures (Park et al. 2009), and microbiome composition (Odamaki et al. 2016). In most cases only a few, most often the first, PCA scores were associated with age and hence could be naturally interpreted as biomarkers of age (Nakamura and Miyao 2007, Nakamura et al. 1988), see Jia et al. (2016) for a recent review and references therein.

The dimensionality reduction revealed by the PCA suggested that the age-associated changes are effectively controlled by the very few variables, naturally associated with the organism-level, or “macroscopic”, properties, such as responses to stresses, risks of diseases and therefore to the remaining health- and life-span (Fedichev 2018, Podolskiy et al. 2015). It is therefore expected that almost any method using organism state variables to predict chronological age, frailty, diseases or the remaining life expectancy should produce the more or less same predictors, albeit at a different signal to noise ratio. This is why biological age estimates are associated with both mortality (Christiansen et al. 2016, Horvath et al. 2015b, Marioni et al. 2015) and morbidity risks (Horvath and Levine 2015, Horvath et al. 2014a,b, 2015a), as well as with life-shortening lifestyles (Gao et al. 2016) or diseases (Horvath and Levine 2015, Horvath et al. 2014a, 2015a, Zhang et al. 2016).

We performed a systematic evaluation of popular biological age models in the same cohort of NHANES study participants. We used a range of machine-learning tools spanning from popular linear chronological age predictors or log-linear models of mortality or morbidity, on one hand, to their fancier deep learning counterparts, on the other. Since the biological age is expected to reflect on the organism’s state rather than on the performance of any specific functional subsystem, we used two sources of the biological information. The first choice was the most commonly used form of clinical blood analysis. In an independent calculation, we used a brief questionnaire developed to gauge the self-reported health of an individual over the last 30 days.

Interestingly, the biological age marker built using blood samples did not dramatically outperform the one obtained from the survey questionnaire. We show that biological aging acceleration of both models is associated with disease burden in persons with diagnosed chronic age-related diseases. For healthy individuals, the same quantity was associated with lifestyles, such as smoking, or variations in the current physical and mental health status. The biological age thus emerged as a universal biomarker of age, frailty and stress, perfectly suitable for applications involving large scale studies of effects of longevity drugs on risks of diseases and on quality of life.

### The limits of deep learning of chronological age

A good biological age predictor should be strongly associated with chronological age and predict the incidence of future diseases and death. We chose these criteria to compare the models using data on lifestyle and chronic health conditions from NHANES (12734 female and 12007 male participants aged 18 – 85). All the study participants were split into training and test cohorts of equal size of 12370 and 12371 subjects. The training set was used to build models and all the results reported below were produced using the test set samples only. We used the same set of log-scaled complete blood cell counts (CBC) and blood biochemistry markers, such as concentrations of C-reactive protein, albumin, alkaline phosphatase, gamma-glutamyl transferase, globulin and serum glucose.

We started by training the baseline biological age model, the “LIN-bioage”, which was a linear regression of the blood features trained to predict the chronological age. The blood markers captured the age-dependence of the organism state fairly well (the Pearson’s correlation of “LIN-bioage” predictor with its target, the chronological age, was *r* = 0.52 with the root mean squared error (RMSE) of 15.9 years). The biological aging acceleration (BAA), i.e. the difference between the predicted biological age of an individual and the mean biological age of their age- and sex-matched peers (Pyrkov et al. 2018a), was associated with the risks of death and hence the remaining life expectancy, see Table I. The effect size and the significance of the association between the BAA and mortality was formally established with the help of the Cox proportional hazards model (Cox 1992) using the subject’s age, sex, smoking and disease (diabetes) status as additional covariates.

**TABLE I:** The association of biological aging acceleration with all-cause mortality, health status and smoking lifestyle in bioage models for the NHANES cohort.

| Covariate | all-cause mortality |  | current/ never smoker |  | health status |  |
| --- | --- | --- | --- | --- | --- | --- |
| | HR (95% CI) | p-value | $\Delta$ bioage | p-value | $\Delta$ bioage | p-value |
| age | 1.09 (1.09 - 1.09) | 7.3E-267 |  |  |  |  |
| sex | 1.57 (1.49 - 1.66) | 1.3E-18 |  |  |  |  |
| diabetes | 1.76 (1.67 - 1.87) | 5.6E-23 |  |  |  |  |
| smoking | 2.35 (2.21 - 2.50) | 6.5E-43 |  |  |  |  |
| + aging acceleration |  |  |  |  |  |  |
| LIN-bioage | 1.04 (1.04 - 1.05) | 9.0E-72 | 2.4 | 6.5E-07 | 2.5 | 1.2E-36 |
| DNN-bioage | 1.05 (1.04 - 1.05) | 1.1E-27 | 1.8 | 1.4E-05 | 0.3 | 2.4E-19 |
| MORBID-bioage | 1.05 (1.04 - 1.05) | 2.7E-72 | 2.1 | 1.8E-09 | 3.0 | 4.2E-55 |
| MORTAL-bioage | 1.08 (1.08 - 1.09) | 4.5E-138 | 2.0 | 3.7E-10 | 0.8 | 2.4E-15 |
| MORTAL-bioage with explicit age | 1.06 (1.06 - 1.07) | 1.2E-163 | 2.6 | 1.1E-13 | 1.8 | 6.4E-28 |
| HAZARDS-Survey | 1.07 (1.06 - 1.07) | 7.2E-77 | 2.6 | 7.0E-15 | 1.8 | 8.8E-32 |
| HAZARDS-Blood-Survey | 1.07 (1.07 - 1.08) | 3.9E-194 | 4.1 | 1.0E-18 | 2.4 | 3.8E-37 |
| Deep HAZARDS-Blood-Survey | 1.10 (1.10 - 1.10) | 5.5E-219 | 4.2 | 6.7E-19 | 2.4 | 2.9E-36 |

Deep neural networks (DNN) can be used to improve age-predicting accuracy by accounting for possible non-linear associations among features. We introduced a “DNN-bioage” predictor, the result of the model trained to minimize MSE between predicted and actual chronological age. The DNN architecture comprised six fully-connected layers, each having from 4 to 12 “leaky ReLU” neurons. The respective “DNN-bioage” model produced a better correlation with age (Pearson’s *r* = 0.64, *RMSE* = 14.6 years) and yet did not demonstrate any improvement of the association with all-cause mortality, see Table I.

We expected the BAA to increase depending on the health status, i.e. in groups of individuals diagnosed with at least one chronic age-related health conditions. More specifically, we formed a list of diseases, including diabetes, high blood pressure, angina pectoris, coronary heart disease, heart attack, heart failure, stroke, arthritis, chronic bronchitis, emphysema and any kind of cancer. The incidence of all the selected health conditions had a strong association with age and increased exponentially with age in the NHANES population. The BAA was significantly associated with the prevalence of the chronic diseases (see Table I). The effect was larger and statistically stronger in the “LIN-bioage” model (2.5 years, *p* = 1.2*E−*36) than in the “DNN-bioage” (0.3 years, *p* = 2.4*E−*19).

In healthy subjects (i.e. those with no diagnosed chronic diseases), both models demonstrated increased biological age in groups of current smokers vs. those who never smoked. Again, the bioage difference was larger and more statistically significant for the “LIN-bioage” model (2.4 years, *p* = 6.5*E −*07) than for “DNN-bioage” (1.8 years, *p* = 1.4*E −* 05).

### Morbidity and mortality models produce the most accurate biomarkers of age

The association of BAA with mortality is the most desired property of a good biological age model. Hence, we turned to log-linear mortality models trained in the same blood markers. We used the NHANES linked death register data (1999–2010 surveys, 7244 female and 7156 male participants aged 40–85, 1517 recorded death events during follow-up till 2011). The first choice was the Cox proportional hazards model with blood markers as covariates only, with no explicit age or sex labels. The log-scaled hazards ratio is a linear combination of blood features and can be calibrated in years as outlined in (Liu et al. 2018, Pyrkov et al. 2018b) and was used as an alternative biological age predictor, the “MORTAL-bioage”. Although the correlation of the predicted age with the chronological age was lower (*r* = 0.35 with *RMSE* = 17.5 years), its association with mortality was notably improved (*HR* = 1.08, *p* = 4.5*E −* 138) relative to that of the models based on the chronological age only, see Table I.

The Cox proportional hazards model relies on the availability of morbidity or mortality follow-up data, which is almost always limited because it requires a long time to collect. Fortunately, the log-hazard ratio of a risk model can be well approximated by a log-odds ratio of logistic regression trained to predict the outcome label whenever the number of the events is sufficiently small (Abbott 1985, Green and Symons 1983). Accordingly, we used the study participant’s health status, i.e. the presence of at least one of the age-related diseases, as the binary morbidity label for the logistic regression model. The resulting log-odds ratio was re-calibrated in years of life and referred to as the “MORBID-bioage” (Pearson’s correlation coefficient to chronological age *r* = 0.48, *RMSE* = 16.2 years). The BAA of both “MORBID-bioage” and “MORTAL-bioage” were significantly associated with all-cause mortality and outperformed the chronological age predictors, see Table I.

The “MORTAL-bioage” model performance could, in principle, be further improved by training a proportional hazard model including the age and sex covariates explicitly. We produced such model and after re-scaling to units of years referred to its log-proportional hazard ratio as the biological age of the “MORTAL-bioage with explicit age” model. The BAA of “MORTAL-bioage” trained with and without explicit age were highly correlated across the samples (Pearson’s *r* = 0.90). Nevertheless, “MORTAL-bioage with explicit age” yielded a slight improvement in its statistical power, see Table I.

In healthy (i.e. chronic disease-free) subjects, the mortality (with and without explicit age) and morbidity models demonstrated an increased and more statistically significant acceleration of bioage for current smokers over non-smokers than that produced by the best chronological age predictor, see Table I. The mortality model with explicit age performed best. Notably, the effect of smoking on the BAA was reversible: there were no significant differences in the bioage levels in groups of never-smokers and those who quit smoking earlier in life, see Fig. 1A.

**FIG. 1:**
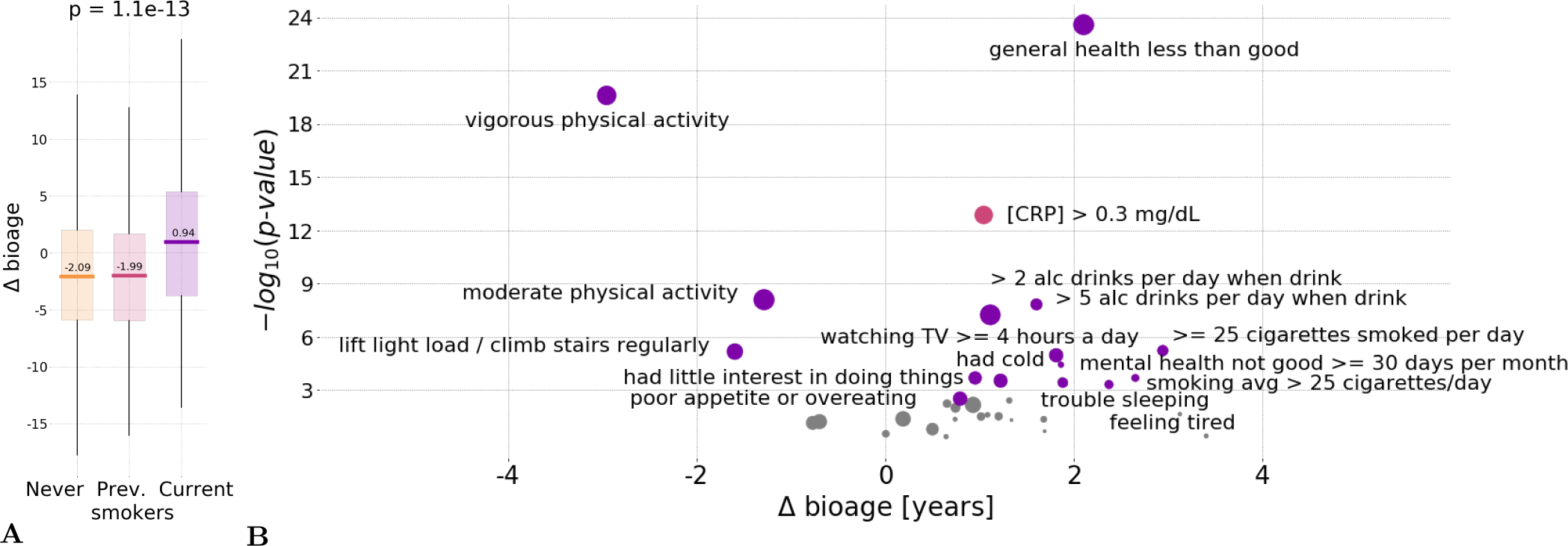
Associations of biological aging acceleration with survey questionnaire responses. **A.** BAA was elevated in groups of current smokers in age- and sex-matched cohorts in NHANES population. The effect of smoking was reversible according to all bioage models, since there were no significant differences between those who never smoked or quit smoking earlier in life (the box-plots here are produced with the help of the “MORTAL-bioage with explicit age” model). **B.** Concentration of C-reactive protein was significantly associated with BAA of the survey-based risk model, the “HAZARDS-Survey” (the red dot). All the other data points (colored in magenta) represent the association effect (the horizontal axis) between the survey responses and the BAA of the “MORTAL-bioage with explicit age” risk model built using the blood analysis variables. Circle sizes correspond to relative number of persons in the smallest of the two compared groups, the largest size corresponds to 50%, i.e. to the groups of equal size.

### Biological aging acceleration, lifestyles, and self-reported physical and mental health

We reasoned that the biological age is an organism-level property, characterizing the overall health of the organism and its interaction with environmental factors. To reveal this connection in detail, we used the BAA of the “MORTAL-bioage with explicit age” model and screened it for associations with self-reported lifestyles and physical and mental health status from the NHANES questionnaire.

In healthy subjects, the BAA was associated with lifestyle characteristics, such as vigorous physical activity (reduced biological age by *≈*3 years) and self-reported general health (increased biological age by *≈*2 years when fair or poor), see Fig. 1B. Smoking had the largest negative effect on biological age which increased by *≈*3 years when smoking more than a pack per day.

Most notably, we observed that several characteristics from the “Depression screener” category also showed association with increased biological age, including self-reported sleeping troubles and feeling tired or having little interest in doing things (*p ≈* 3*E −* 4).

### Biological age based on self-reported well-being questionnaire

So far, we focused on comparing the performance of biological age predictor’s association with mortality and morbidity depending on various machine learning approaches involved in the training of the models and on different learning targets, such as chronological age or age at death. In this section we turn to investigation of the role of the source of biological signal. The use of clinical blood analysis variables is common, and yet requires an invasive medical procedure. Instead, we proposed using a brief questionnaire, including a dozen of questions related to overall physical and mental health of an individual over the one month prior to the assessment.

We used various data fields of the NHANES survey database with the best associations with all-cause mortality and characteristic of the current health status of a study participant on a relatively short period of less than a year, most during the last three months or less, see Table II. The data-fields marked with a star (“*”) were available for all studied NHANES cohorts (1999–2010) and were used as covariates in hazards models. We built another Cox proportional hazards model and transformed its log-hazard ratio into the “HAZARDS-Survey” bioage, using the selected set of questionnaire data fields including chronological age and sex as covariates. The resulting predictor demonstrated a good association of its BAA with all-cause mortality at the significance level of *p* = 7.2*E −* 77, see Table I. Somewhat surprisingly, the statistical power of the association of the BAA of the “HAZARDS-Survey” was poorer, although not dramatically worse than that of “MORTAL-bioage with explicit age” model, which was trained in the clinical blood markers and explicit age.

**TABLE II:** NHANES survey data fields.

| NHANES Variable code | NHANES Variable name (as in 2005-2006 cohort) |
| --- | --- |
| HSD010 | General health condition (*) |
| HSQ470 | no. of days physical health was not good (*) |
| HSQ480 | no. of days mental health was not good (*) |
| HSQ490 | inactive days due to phys./mental hlth (*) |
| HSQ500 | SP have head cold or chest cold (*) |
| HSQ510 | SP have stomach or intestinal illness? (*) |
| HSQ520 | SP have flu, pneumonia, ear infection? (*) |
| ALQ130 | Avg no. alcoholic drinks/day -past 12 mos (*) |
| WHQ070 | Tried to lose weight in past year (*) |
| WHQ090 | Tried not to gain weight in past year (*) |
| PAD020 | Walked or bicycled over past 30 days (*) |
| PAQ180 | Avg level of physical activity each day (*) |
| PAD590 | no. hours watch TV or videos past 30 days (*) |
| PAD200 | Vigorous activity over past 30 days (*) |
| PAD320 | Moderate activity over past 30 days (*) |
| PAD460 | Number of times past 30 days (*) |
| RDQ031 | Coughing most days - over 3 mo period (*) |
| RDQ050 | Bring up phlegm most days - 3 mo period (*) |
| RDQ070 | Wheezing or whistling in chest - past yr (*) |
| RDQ135 | Limit usual activities due to wheezing (*) |
| RDQ140 | Had dry cough at night in past year (*) |
| DPQ010 | Have little interest in doing things |
| DPQ020 | Feeling down, depressed, or hopeless |
| DPQ030 | Trouble sleeping or sleeping too much |
| DPQ040 | Feeling tired or having little energy |
| DPQ050 | Poor appetite or overeating |
| DPQ060 | Feeling bad about yourself |
| DPQ070 | Trouble concentrating on things |
| DPQ080 | Moving or speaking slowly or too fast |
| DPQ090 | Thought you would be better off dead |
| DPQ100 | Difficulty these problems have caused |
| SMD070 | no. cigarettes smoked per day now |
| SMD641 | no. days smoked cigs during past 30 days |
| SMD650 | Avg no. cigarettes/day during past 30 days |
| RIDAGEYR | Age (*) |
| RIAGENDR | Gender (*) |

The BAA of the pure blood markers- and the survey-based only models were almost equally well associated with the presence of chronic diseases (*p* = 6.4*E −* 28 and *p* = 8.8*E −* 32, respectively). Both models were sensitive to smoking status and predicted 2.6 years of the bioage difference at the significance level of *p* = 1.1*E −* 13 and *p* = 7.0*E −* 15, respectively (Table I), even though we intentionally excluded any data fields directly related to smoking from the Survey data fields. The questionnaire and the blood markers could be combined and used for training of a more powerful, the combined “HAZARDS-Blood-Survey” bioage model that yielded a minimally improved association with all-cause mortality (*p* = 3.9*E −* 194, see Table I). The combination of the two sources of biological data immediately yielded a significant improvement in association with the health status (2.4 years, *p* = 3.8*E −* 37) and smoking (4.1 years, *p* = 1.0*E −* 18).

The BAA of the risk models based on the questionnaire- and clinical blood analysis were significantly associated with the levels of C-reactive protein (CRP, see Fig. 1B).

### Deep learning of proportional hazards models and biomarkers of aging discovery

Finally, we used the blood markers and questionnaire features combined to train a DNN-based hazards model “Deep HAZARDS-Blood-Survey”. The model was based on the same architecture as “DNN-bioage” with loss function replaced by negative logarithm of Cox-Gompertz likelihood adopted from (Bender et al. 2005):

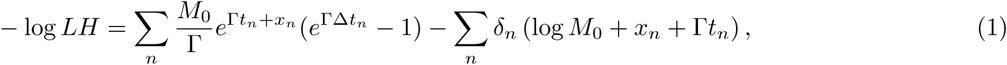

where *M*_0_ and Γ are the initial mortality rate and mortality doubling rate of the Gompertz mortality law, *t*_*n*_ is the age of *n*-th participant, *δ*_*n*_ and Δ*t*_*n*_ are the event outcome and follow-up time till event or censorship, and *x*_*n*_ is the output of DNN. The parameters of the baseline Gompertz mortality model were fitted to NHANES follow-up data without the DNN output (*x*_*n*_ = 0). We obtained *M*_0_ = 5.7*×*10^−5^ and Γ = 0.084 per year corresponding to the average lifespan of 79.9 years (see, e.g., Tarkhov et al. (2017) for the discussion of the Gompertz model). The model estimate was in a good agreement with the life expectancy at birth of 79.3 years reported for the US (WHO 2016).

In limited cohorts of human subjects, the simultaneous determination of small corrections to the Gompertzian variables Γ and *M*_0_ is a poorly defined mathematical problem (Tarkhov et al. 2017). We, therefore, held the baseline model parameters fixed in Eq. 1 and optimized the deep network to include the physiological-state dependent variables through the network output variable *x*_*n*_ (Pyrkov et al. 2018b). As for all other models above, the log-hazard ratio of the deep mortality model was re-scaled to years and referred to as the “Deep HAZARDS-Blood-Survey” bioage.

As expected, the model predictor demonstrated a better association with all-cause mortality. We did not, however, observe any significant improvement of association of the model BAA with the health status (*p* = 2.9*E −* 36) and smoking status in the healthy individuals (*p* = 6.7*E −* 19). Nevertheless, the calculation shows that deep biological age models based on Eq. 1 may be a good starting point for state-of-the-art biomarkers of aging discovery. The suggested procedure explicitly exploits the exponential nature of the mortality acceleration characteristic to aging process and deep learning architectures for automatic non-linear feature extraction.

## Discussion

A good biomarker of aging should predict the remaining health- or lifespan and, at the very best, be causally associated with the underlying biology of aging. We presented a systematic investigation of biological age predictors trained in sets of various physiological indices (clinical blood biomarkers or a self-reported questionnaire). Using chronological age as the target for supervised training of biological age models is almost always a bad choice, since it implies minimization of the discrepancy between the predicted and chronological age (i.e. aging acceleration) and thus effectively destroys the model sensitivity to changes in health. This undesired effect was especially aggravated in combination with deep neural networks (DNN). At the same time, the exponential Gompertz mortality rate acceleration in human populations suggests that proportional mortality and morbidity hazards models can be used as powerful alternatives to obtain a novel biomarker of the organism’s state, reflecting the levels of external and endogenous stresses.

Proportional hazards mortality models are increasingly common tool for biological age predictions. Most recent examples include the “PhenoAge” based on blood sample data (Liu et al. 2018). The “PhenoAge” prediction was further used to train a DNA-methylation-based aging marker DNAm PhenoAge (Levine et al. 2018). The number of samples in both studies was comparable to that in our present work. DNAm PhenoAge was reported to be associated with all-cause mortality at significance level *p* = 9.9*E −* 47 in a meta-study comprising samples from the Framingham Heart Study, the Normative Aging Study, and the Jackson Heart Study (*≈*9000 participants with *≈*2000 death events observed in follow-up). The authors also mentioned that its performance was superior to epigenetic biomarkers of aging (see, e.g., the value of *p* = 1.7*E −* 21 reported in (Levine et al. 2018) for Hannum model (Hannum et al. 2013)) trained to predict chronological age. This result also seems consistent with with our findings here with clinical blood markers and earlier in analysis of human physical activity-based bioage models (Pyrkov et al. 2018b). The overall conclusion is that the most accurate chronological age predictors produced the poorest associations with all-cause mortality (see Table I) and thus should be avoided whenever possible in favor of the explicit mortality or morbidity models.

Advanced machine learning tools are naturally called to improve the biological age predictions, see, e.g., a model trained from clinical blood markers (Putin et al. 2016) and facial photos (Bobrov et al. 2018). The examples presented here show that the full power of the deep learning architectures could be harnessed for feature extraction and non-linear models fitting of risks, rather than chronological age models (Pyrkov et al. 2018b). The risk models, however, require follow-up information involving the incidence of age-related diseases or death. The exponential nature of mortality and morbidity acceleration implies that a risk model could be approximated by a logistic regression (Abbott 1985, Green and Symons 1983, Pyrkov et al. 2018a) to health status. This is especially useful, since morbidity data is easier to collect. We established a high concordance and comparable statistical power of the models trained to predict mortality and morbidity risks or prevalence of chronic diseases.

Training a chronological age prediction model from biological signals is a common approach to produce a biological age model, such as e.g., gene expression (Peters et al. 2015), IgG glycosylation (Kristic et al. 2014), blood biochemical parameters (Levine 2013), gut microbiota composition (Odamaki et al. 2016), and cerebrospinal fluid proteome (Baird et al. 2012). Some physiological indices are better correlated to age. For example, the Pearson’s correlation of our “LIN-bioage” biological age predictor with age was only *r* = 0.52, that is significantly lower than *r* = 0.65 – 0.70 reported for models based on IgG-glycosylation and *r ≈* 0.90 for proteome (Enroth et al. 2015) or DNA-methylation data (Hannum et al. 2013, Horvath 2013). The profound correlations of specific physiological indices with the chronological age may still be an important signature of the organism state dynamics not directly associated with stress, morbidity and mortality.

As far as we can judge from a comparison of methods for assessing mortality risks (Levine and Crimmins 2014), the bioage model produced in DNA-methylation data and modeling of mortality risks had a similar association with mortality and health risks as those built in the present work using blood sample and survey questionnaire. DNAm PhenoAge was reported as *≈*2 years different in cohorts of smokers and non-smokers at the significance level *p* = 0.0033 in a group of 209 current, 701 former and 1097 never-smokers (Levine et al. 2018). Our models produced the bioage differences from 2 to 4.2 years in cohorts including 530 current and 1201 never-smokers in the test set at the significance level ranging from *p ≈* 1.0*E* − 5 to 6.7*E −* 19, depending on the model. The biological aging acceleration in the DNAm PhenoAge and in all our models was smaller, than the expected lifespan difference between smokers and non-smokers (which was estimated to be as much as ten years (Doll et al. 2004)). The statistical power of, say, the “MORTAL-bioage” with explicit age model would drop to comparable to that of DNAm PhenoAge if the number of samples would roughly be halved to match the cohort sizes of (Levine et al. 2018).

In agreement with our findings here and in the technically similar analysis of physical activity data (Pyrkov et al. 2018a), the BAA of the DNAm Phenoage was associated with smoking and, in the same time, was not different in groups of never smokers and those who quit smoking early in life. We note, that the BAA remained significantly associated with smoking even if we restrict ourselves to the chronic disease-free subjects only and hence could not be simplified only to the excess of the disease burden inflicted by smoking. Another notable BAA association in the healthy cohorts was that with CRP. The latter is a molecular marker of inflammation caused by a wide variety of conditions, from infections to cancer (Heikkilä et al. 2007). We, therefore, argue that the BAA is a universal stress indicator, a signature of the organism-level stress response to a generic environmental or endogenous factors, including lifestyles and diseases.

Similar performance of all hazards-based bioage models regardless of the source of biological signal suggests that all of them extract the same underlying biological factor, which manifests itself on the organism level and could be measured with similar accuracy in biological signals of different kind. The biological aging acceleration of these models is associated with disease burden in persons with diagnosed chronic age-related diseases. For healthy individuals, the same quantity was associated with smoking, current physical and mental health. The biological age thus emerges as a universal biomarker of age, frailty and response to stress suitable for applications involving large scale studies of the effects of future anti-aging drugs and life-style interventions on risks of diseases and on quality of life.

## Acknowledgements

The authors would like to thank Konstantin Avchaciov from Gero team for proof reading and thoughtful comments. The work was suppored by Gero LLC.

